# Misophonia Symptoms Severity is Attributed to Impaired Flexibility and Heightened Rumination

**DOI:** 10.1101/2025.01.13.632542

**Authors:** Vivien Black, J.D. Allen, Hashir Aazh, Sheri L. Johnson, Mercede Erfanian

**Affiliations:** Department of Psychology, University of California, Berkeley, CA, USA; Hashir International Specialist Clinics & Research Institute for Misophonia, Tinnitus and Hyperacusis, London, United Kingdom; ESSCA School of Management, Lyon, France

**Keywords:** Misophonia, affective flexibility, cognitive flexibility, rumination

## Abstract

Misophonia is a disorder involving sensitivity to certain sounds and related stimuli. Here, we explore the relationship between misophonia and affective flexibility, which describes cognitive shifting abilities in the face of emotion-evoking stimuli. Previous evidence suggests impaired subjective cognitive flexibility in misophonia, but this relationship has not been confirmed behaviourally or in emotionally-salient contexts. The secondary aim of this study is to test the potential association between misophonia and cognitive flexibility, building upon findings from previous research. The third objective is to examine the relationship between misophonia and rumination, a maladaptive cognitive process characterized by repetitive negative thinking and linked to both cognitive and affective inflexibility. One hundred and forty participants completed the recently developed Memory and Affective Flexibility Task (MAFT), designed to assess affective flexibility, as well as a battery of self-report measures to evaluate misophonia severity, cognitive flexibility and rumination. Results suggested an inverse relationship between affective flexibility as measured by switch accuracy, but not reaction time, and misophonia severity. Cognitive flexibility was also inversely associated with misophonia severity, but notably did *not* attribute to task-based affective flexibility, suggesting two independent constructs both involved in misophonia manifestation. Rumination associated positively with misophonia severity and inversely with cognitive flexibility, but not affective flexibility. Taken together, these findings highlight a unique cognitive profile of misophonia, characterized by rigidity at the psychological level through cognitive inflexibility and rumination, as well as at the executive-function level in terms of affective switching difficulties.

## Introduction

Misophonia is a disorder characterized by decreased tolerance to certain sounds, such as orofacial (e.g., chewing or sniffing), non-orofacial but human-based (e.g., knuckle cracking), or innocuous environmental sounds (e.g., clock ticking). Among individuals with misophonia, these trigger sounds, and sometimes also visual (misokinesia) or other related sensory stimuli, evoke strong emotional reactions such as anger, irritation, disgust, and (anticipatory) anxiety (Swedo et al., 2022). These emotional reactions are accompanied by autonomic nervous system arousal (Edelstein et al., 2013; Oszczapinska et al., 2024) and altered neural activity in areas including those related to emotion processing, the salience of sensory stimuli, and motor control (Kumar et al., 2021; Kumar et al., 2017; Schroder et al., 2019). Resulting behavioural responses, such as aggression, anger outbursts and avoidance, often cause significant impairment in daily life, especially in the case of more severe misophonia (Brout et al., 2018). Despite prevalence estimates ranging from 4.6% to 20% (Dixon et al., 2024; Vitoratou et al., 2023; Wu et al., 2014; Zhou et al., 2017), research on misophonia, especially its treatment, is still at its early stages (Rosenthal et al., 2023; Swedo et al., 2022)

Evidence has highlighted the presence of executive functioning deficits in individuals with misophonia in areas such as cognitive control, response bias, emotion regulation and attentional processing (Daniels et al., 2020; Eijsker et al., 2019; Guetta et al., 2022; Murphy et al., 2024; Simner et al., 2021). Those affected often show a tendency to hyper-focus on certain bothersome sounds (misophonic triggers) and face significant challenges in shifting attention away from these stimuli (Edelstein et al., 2013; Jager et al., 2020). They show selective attention deficits in response to trigger sounds as well as hypervigilant anticipation, including anticipatory anxiety (Murphy et al., 2024; Sanchez & Silva, 2018). This pattern may reflect deficits in attentional set-shifting, a core feature of cognitive inflexibility, which is defined as the inability to disengage from irrelevant stimuli and redirect attention to information relevant to the task or situation at hand (Horning & Davis, 2012; Uddin, 2021). In support of this idea, there is evidence of higher self-reported cognitive inflexibility among those with misophonia (Simner et al., 2021). Cognitive flexibility, which facilitates efficient shifting between mental processes to produce contextually appropriate behavioural responses, follows a protracted developmental trajectory and is frequently impaired in numerous common neurodevelopmental and psychiatric disorders (Dajani & Uddin, 2015; Morris & Mansell, 2018), including obsessive-compulsive disorder (OCD) (Gruner & Pittenger, 2017), anorexia nervosa (AN) (Miles et al., 2020), autism spectrum disorder (ASD) (Albein-Urios et al., 2018; Lage et al., 2023), Post-Traumatic Stress Disorder (PTSD) (Popescu et al., 2023), and attention-deficit and hyperactivity disorder (ADHD) (Zhang et al., 2023). Misophonia shows substantive comorbidity with these conditions, and the presence of such comorbidities relate to severity of misophonia as well (Andermane, Bauer, Simner, et al., 2023; Erfanian et al., 2019; Kluckow et al., 2014; Rinaldi et al., 2023; Rouw & Erfanian, 2018; Wu et al., 2014). The occurrence of these symptoms suggests potential overlap of neurobiological or psychological mechanisms, and cognitive flexibility is one such candidate mechanism.

It remains, however, unclear whether cognitive inflexibility in misophonia manifests primarily in the face of triggers vs. across a broad range of contexts. There are some indications that the cognitive inflexibility may be broad. For instance, misophonia symptoms are associated with a greater aversion to expectation violation in social contexts, suggesting a broader inflexible application of social standards to others’ behaviour (Banker et al., 2022). Furthermore, misophonia has repeatedly been associated with perfectionism (Jager et al., 2020; Jakubovski et al., 2022), which is a process associated with cognitive inflexibility, characterized by a *rigid* pursuit of unrealistic personal goals and standards (Egan et al., 2011). In other words, perfectionism may contribute to misophonia by fostering a tendency to inflexibly identify with negative self-views when self-imposed standards are not met (Nguyen & Morris, 2024). Perfectionists show an attentional bias that directs greater focus toward negative information than positive information (Shafran et al., 2002). In the context of misophonia, this translates to an intensified focus on negatively perceived sounds (Simner et al., 2021). These findings may reflect cognitive inflexibility among individuals with misophonia that extends beyond contexts specifically associated with triggering stimuli.

### Task-Based correlates of cognitive flexibility in misophonia

To understand cognitive flexibility in misophonia, it is crucial to examine whether deficits shown on self-report data can also be reflected in a neurocognitive task. Self-report data may not accurately capture deficits in executive functioning as they tend to emphasize personality-level traits of flexibility rather than task-specific performance (Buchanan, 2016). In this respect, behavioural studies have yet to establish a consistent link between task-based cognitive flexibility and misophonia. For instance, despite reduced cognitive flexibility reported in questionnaires (Simner et al., 2021), individuals with misophonia do not appear to perform differently on the Wisconsin Card Sorting Task (WCST) (Abramovitch et al., 2023), a task in which perseverative responses/perseverative errors are often considered to reflect cognitive inflexibility (Miles et al., 2021). This discrepancy may stem from inconsistent definitions of cognitive inflexibility, as well as methodological and reporting biases, which likely underlie lack of connection between self-reported measures of cognitive flexibility and performance on neuropsychological tasks. In addition, self-report measures may not be reliable proxies for neuropsychological tasks of cognitive flexibility (Howlett et al., 2021). As such, the self-reported cognitive flexibility among those with misophonia (Simner et al., 2021) may not inform the extent to which individuals with misophonia show impairments in set shifting or task switching, core components intricately linked to cognitive flexibility (Dajani & Uddin, 2015; Stemme et al., 2007).

### Affective Flexibility in Misophonia: Cognitive Flexibility in Emotional Contexts

Another possible reason for the previous null findings regarding task-based cognitive flexibility in individuals affected by misophonia, would be that cognitive inflexibility is evoked only in the presence of triggers (e.g., orofacial sounds) or other relevant stimuli. For instance, the association between impaired cognitive control and misophonia severity was specific to performance on the Stroop task in the presence of misophonia trigger sounds, with misophonia severity not relating to Stroop effect when faced with sounds considered to be generally unpleasant (Daniels et al., 2020). This could indicate that specific executive functioning deficits may only be present in certain contexts among individuals with misophonia.

Further, the WCST contains only neutral stimuli, which we hypothesize may not be sufficient to evoke impairments in shifting or switching abilities in misophonic people. Considering the role of emotion regulation deficits in misophonia (Dixon et al., 2024; Guetta et al., 2022), emotionally evocative stimuli could be necessary to evoke significant inflexibility in behavioural responding during a task by individuals with misophonia. The potential importance of affective stimuli to evoking flexibility deficits can be shown by drawing parallels with previous research in ASD, which has shown a relationship with misophonia in the form of heightened presence of autistic traits (Rinaldi et al., 2023). Task-switching paradigms revealed that switch costs associated with autism were specific to trials involving switching attention either to or away from emotional stimuli, with directional findings referring to variations in these costs based on the direction of the shift. In other words, these costs differ by direction, reflecting distinct cognitive processes engaged by emotional salience (De Vries & Geurts, 2012; Latinus et al., 2019). Cognitive flexibility in response to emotion-evoking stimuli is referred to as *affective flexibility* (Eckart et al., 2021). Hence in the present study, we use a novel neurocognitive task, the Memory and Affective Flexibility Task (Allen et al., Under review) that incorporates neutral and emotionally-arousing stimuli in working memory and affective flexibility trials (Lang & Bradley, 2007).

### Rumination: A key process related to inflexibility potentially involved in misophonia

Rumination is recognized as a transdiagnostic process associated with cognitive and affective inflexibility (Altamirano et al., 2010; Davis & Nolen-Hoeksema, 2000; Genet et al., 2013; Morris & Mansell, 2018). Defined as repetitive, self-focused thought related to negative emotions and their origins, rumination is pervasive across a range of psychiatric disorders (Treynor et al., 2003). Inherently rigid and maladaptive, rumination involves cycling through the same negative thoughts without transitioning to more adaptive cognitive approaches. Inflexibility may manifest in rumination through deficits in the ability to disengage from negative emotional elements of stimuli, resulting in difficulty shifting focus from negative to neutral or positive cognitions (Genet et al., 2013; Koster et al., 2011). In misophonia, rumination may manifest as inflexible, repetitive negative thinking, not only about trigger stimuli and their sources but also involving internally focused thoughts, such as judgments about the misophonic reaction itself. Misophonic individuals tend to ruminate even when no triggers are present, such as by anticipating what trigger sounds may come next or replaying previous triggering situations (Vitoratou et al., 2021). Although people with misophonia do anecdotally report rumination surrounding their misophonic experience, it remains unclear to what extent ruminative thinking is pervasive in broader contexts as an overarching cognitive tendency in people with misophonia. Thus, In the current investigation, we ask whether this tendency to ruminate extends more generally to daily life, both about misophonia and other unrelated topics. We aim to explore rumination as a potential manifestation of cognitive and affective inflexibility in misophonia, focusing here on three subtypes: perseverative thinking (Ehring et al., 2011), brooding (Treynor et al., 2003), and anger rumination (Sukhodolsky et al., 2001).

### Current study

By employing a neurocognitive task (Allen et al., Under review), the first aim is to evaluate the relationship between affective flexibility performance and the severity of misophonia symptoms. The MAFT is designed to assess affective flexibility, which may be relevant to individuals with misophonia as well as broader psychopathology (Friedman & Banich, 2019). We predict an association of self-reported misophonia severity with affective inflexibility, as indexed by worse performance on the MAFT switch trials in terms of reduced accuracy and slower Reaction Time (RT) (H1).

In addition, we examine the relationship between self-reported cognitive inflexibility and the severity of misophonia, expecting to replicate the findings of Simner et al. (H2) and we predict a positive association between diminished cognitive flexibility and the severity of misophonia symptoms. Furthermore, we evaluate the association between self-reported cognitive inflexibility and task-based indices of affective flexibility, specifically switch accuracy and switch RT, derived from the MAFT to examine whether self-report flexibility aligns with task-based measures. We hypothesize that self-reported and task-based measures of flexibility will be positively correlated, showing a convergence between cognitive and affective dimensions of flexibility (H3). This investigation aims to deepen understanding of the interplay between subjective and objective facets of flexibility and their relevance to misophonia severity.

Additionally, we explore the association between rumination, misophonia severity, affective flexibility, given affective inflexibility, specifically the difficulty in disengaging from processing the emotional aspect of negative stimuli, has been linked to increased rumination in daily life (Genet et al., 2013). In particular, we expect to find a positive relationship between increased self-reported rumination, and misophonia severity as well as worse switch accuracy and switch RT to correspond with greater levels of rumination (H4). As part of our exploratory analyses, we also examine the relationship between cognitive flexibility and measures of rumination. We hypothesize that decreased cognitive flexibility will be positively correlated with increased levels of rumination (H5).

## Method

Ethical approval for the study was obtained from the Committee for Protection of Human Subjects at the University of California, Berkeley (CPHS protocol ID 2024-01-17038). The study was pre-registered on OSF (https://doi.org/10.17605/OSF.IO/UCBG4) before the creation of data.

### Participants

The sample comprised 140 participants (mean age = 29.98 ± 6.72 years; 49 females, 91 males). Of those, 128 participants were recruited from Prolific (www.prolific.com), an online study participant pool, and 12 were volunteer participants recruited from misophonia-related newsletters; specifically, the soQuiet Misophonia Research Pool (www.soquiet.org/pool) and Misophonia International (www.misophoniainternational.com).

Based on the recommended cut-off score of 87 or above on the S-Five (Vitoratou et al., 2023), 25% of the total sample (*N* = 35) experienced clinically significant misophonia (**Table 1**). Specifically, 18.75% of Prolific participants (*N* = 24) endorsed significant misophonia. This rate is highly similar to the rate of significant misophonia on the S-Five (18.4%) found by previous research in a general population (Vitoratou et al., 2023). Out of the volunteers identified from misophonia-related newsletters, 91.67% (*N* = 11) endorsed significant misophonia (**Figure 1**).

**Figure 1.**
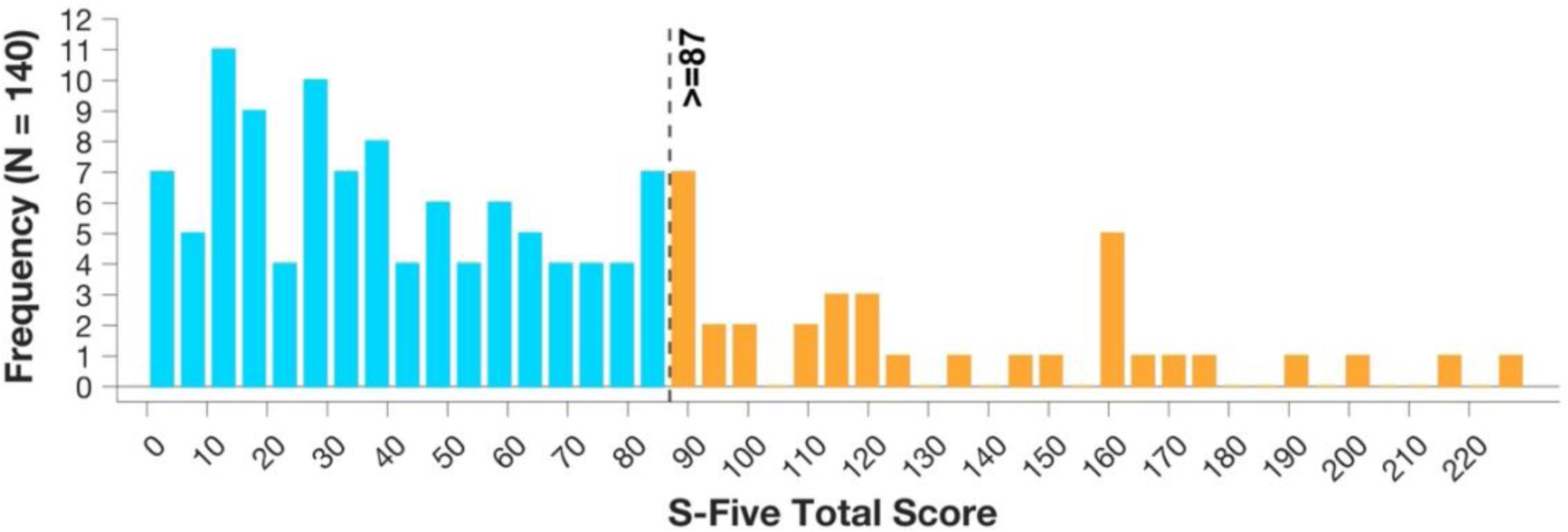
Histogram of S-Five total score distribution. The histogram demonstrates the distribution of total scores on the S-Five Scale assessing misophonia severity. The dashed vertical line represents the threshold for significant misophonia (87 or above) as indicated by (Vitoratou et al., 2023). 25% of the sample (N = 35) experienced significant misophonia based on this measure. Scores ranged between 0 and 226 (S-Five maximum possible score is 250).

**Table 1.** Table below shows between-group demographic comparisons of participants with/without ‘significant’ misophonia. ‘Sig-misophonia’ refers to individuals with clinically significant misophonia, whereas ‘non-sig-misophonia’ denotes those without clinically significant symptoms (p ≥ 0.05).

| Parameter | Sig-Misophonia<br>M(SD) / %(n) | Non-sig-Misophonia<br>M(SD) / %(n) | t/$\chi^2$ | p | Cohen’s d |
| --- | --- | --- | --- | --- | --- |
| **Gender** |  |  | **4.62** | **0.03** |  |
| Female | 51.43% (18) | 29.52% (31) |  |  |  |
| Male | 48.57% (17) | 70.48% (74) |  |  |  |
| **Age** | 31.29 (6.91) | 29.54 (6.63) | -1.31 | 0.20 | -0.26 |

### Materials

#### Memory and Affective Flexibility Task (MAFT, (Allen et al., Under review))

First, participants completed the Memory and Affective Flexibility Task (MAFT) via Inquisit Web (www.millisecond.com) version 6.1.1.

In the MAFT, participants engaged in two types of trials: memory trials and emotional switch trials. Memory trials required participants to determine whether the current image matched one shown *N* trials earlier, measuring working memory. Emotional switch trials, randomly interspersed within the task, required participants to rapidly evaluate whether an image was positive or negative, measuring affective flexibility. The working memory trials were based on the widely used *N*-back task (Owen et al., 2005), designed to measure behavioural responses to emotional and non-emotional stimuli. Switch trials assessed affective flexibility, as they required participants to shift from a memory-based response to evaluating emotional valence, reflecting task switching in the context of affective stimuli. For the current study, only switch accuracy and switch RT were of interest (**Figure 2**).

**Figure 2.**
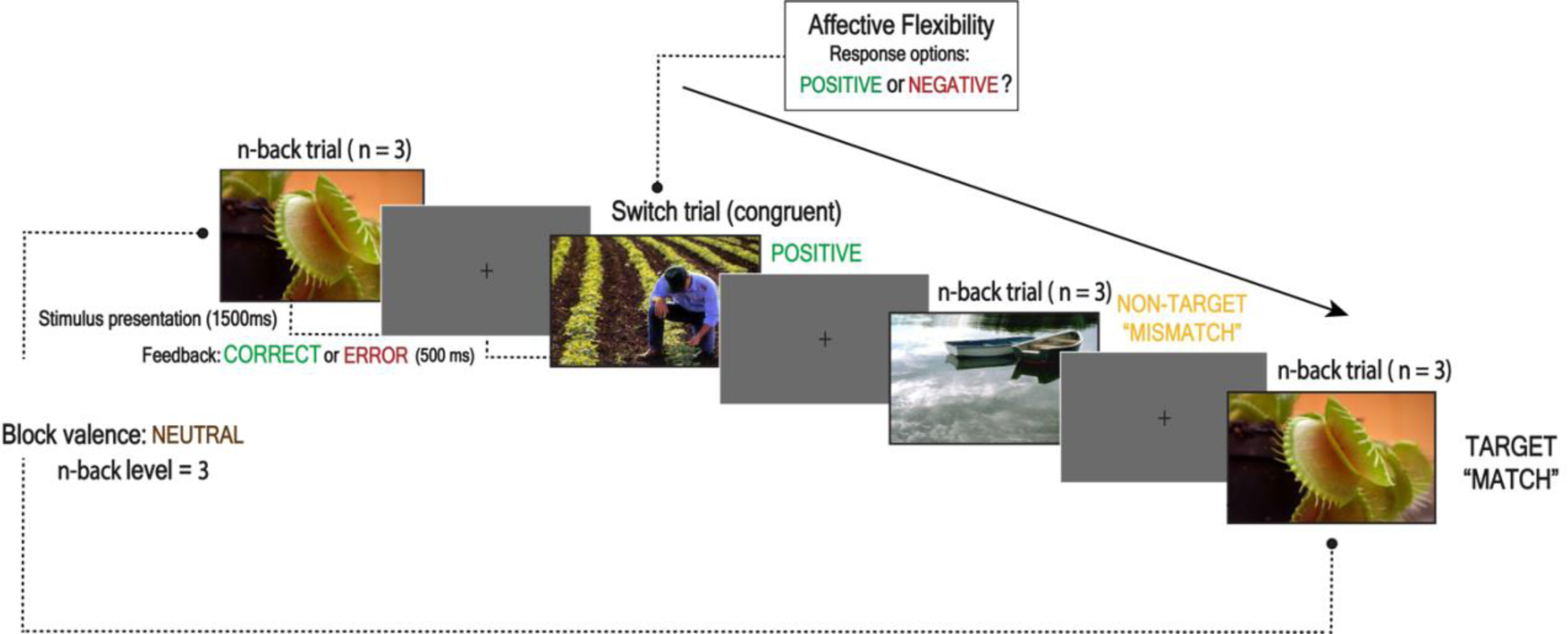
Sequence and timing of stimulus events in the Memory and Affective Flexibility (MAFT) task. A standard block design was used, in which images were presented for 1500 ms with 500 ms fixation cross as intertrial intervals. Participants judged whether the current image matched or mismatched the image presented 1, 2, or 3 trials prior by pressing “A” for match and “L” for mismatch. After each trial, participants received visual feedback indicating the accuracy of their response, displayed as either “CORRECT” or “ERROR” on the screen. In switch trials, they assessed the valence of emotional images (positive, negative or neutral) from the International Affective Picture System (IAPS) (Lang & Bradley, 2007), pressing “K” for negative and “S” for positive.

The MAFT consisted of 198 trials organized into nine experimental blocks, with participants completing three valence-specific blocks (positive, negative, neutral) for each level of *N*-back difficulty (*N* = 1, 2, and 3). Each block included 20 + *N* trials to account for initial “mismatch” stimuli before the first potential “match” target. Prior to the experimental blocks, participants completed two practice blocks with 500 ms trial-level feedback, repeating them until they achieved at least 70% accuracy on *N*-back trials at *N* = 1 (11 trials) and *N* = 2 (12 trials). Approximately 60% of trials within each block were *N*-back trials (12 + *N* per block), consisting of four “match” trials and the remainder as “mismatch” trials, with emotional valences congruent with the block type. The remaining 40% (∼8 trials per block) were switch trials, which alternated between emotional valences and were presented in a pseudo-randomized order. Stimuli were drawn without replacement from the International Affective Picture System (IAPS) image pool (Lang & Bradley, 2007), with negative and positive image sets matched on standardized arousal and valence ratings (negative: arousal M = 5.94 ± 0.77; valence M = 7.22 ± 1.04; positive: arousal M = 5.22 ± 1.02; valence M = 7.15 ± 0.79), and neutral images selected for low arousal (M = 2.88 ± 0.57) and intermediate valence (M = 4.98 ± 0.30) (**Figure 2**).

Performance metrics included accuracy, calculated as the proportion of correct “match” hits and “mismatch” rejections for *N*-back trials and the proportion of correctly classified positive and negative stimuli for switch trials, as well as RT, calculated as the mean response time for correct trials, consistent with prior research (Eckart et al., 2021, Malooly et al., 2013). Neutral stimuli in switch trials were analysed for interpretive bias, with the proportion categorized as negative serving as an index. Neutral stimuli were predominantly classified as positive (M = 0.64 ± 0.11), providing insight into the evaluation of stimuli without salient emotional content.

In memory trials, participants responded by pressing “A” for a match and “L” for a mismatch, while in switch trials, they judged emotional valence, pressing “K” for negative and “S” for positive. Recent findings suggest that RT may not be specific to trial type, as RT indices are highly correlated across tasks, indicating it may reflect general motor or cognitive processing speed rather than task-specific variables (Allen et al., Under review). Although both accuracy and RT were pre-registered for the switch task, recent evidence highlights accuracy as the more meaningful measure of affective flexibility; therefore, both metrics are reported here.

#### Selective Sound Sensitivity Syndrome Scale (S-Five, (Vitoratou et al., 2021))

Misophonia symptom severity was assessed using the 25-item Selective Sound Sensitivity Syndrome Scale (S-Five) (Vitoratou et al., 2021). The S-Five scale is designed to assess five different dimensions of misophonia severity: externalizing, internalizing, impact, threat, and outburst, as well as overall severity. Participants are asked to respond to questions (e.g. “The way I react to certain sounds makes me wonder whether deep inside I am just a bad person” and “The way I feel/react to certain sounds will eventually isolate me and prevent me from doing everyday things”) on a 10-point scale, ranging from (0) for not at all true to (10) for completely true. The maximum possible score on the S-Five is 250. A score of 87 or above indicates significant misophonia as suggested by ROC analysis (Vitoratou et al., 2021). The S-Five has previously exhibited good internal consistency (*α* >= 0.83) and strong test-retest reliability (ICC >= 0.86) for all factors, as well as good convergent and discriminant validity. In the current study, the S-Five showed excellent internal consistency (*α* = 0.96).

#### Detail and Flexibility Questionnaire (Dflex, (Roberts et al., 2011))

The 12-item flexibility subscale of the Detail and Flexibility Questionnaire (DFlex) was included as a self-report measure of cognitive inflexibility (Roberts et al., 2011). In a previous study, participants classified as having misophonia scored significantly higher for inflexibility on this questionnaire compared to those without misophonia (Simner et al., 2021). Participants rated each item (e.g., “I can be called stubborn or single minded as it is difficult to shift from one point of view to another”) on a 6-point Likert scale ranging from Strongly Disagree (1) to Strongly Agree (6). Total scores range from 12 to 72, with higher scores indicating greater levels of inflexibility. This scale has previously demonstrated very high internal consistency (*α* = 0.93) as well as strong construct and discriminant validity. In the current study, the DFlex flexibility subscale showed good internal consistency (*α* = 0.83).

#### The Perseverative Thinking Questionnaire (PTQ, (Ehring et al., 2011)

Perseverative thinking, a measure of Repetitive Negative Thinking (RNT), was assessed using the Perseverative Thinking Questionnaire (PTQ) (Ehring et al., 2011). RNT describes the transdiagnostic process of unproductive, repetitive thinking, and rumination is one form of RNT (Ehring et al., 2011). Participants were asked to rate how each of 15 statements applied to them in terms of how they typically think about their negative experiences or problems. Participants responded using a 5-point Likert scale, from Never (0) to Almost Always (4). Example statements include “The same thoughts keep going through my mind again and again” and “I can’t do anything else while thinking about my problems”. The higher-order PTQ scale has previously exhibited excellent internal consistency (*α* = 0.94 - 0.95), as well as good internal consistency for subscales (Core Characteristics of RNT: *α* = 0.92 - 0.94; Unproductiveness of RNT: *α* = 0.77 - 0.87; RNT Capturing Mental Capacity: *α* = 0.82 - 0.90); satisfactory test-retest reliability (total score retest correlation = 0.69), and good convergent validity (Ehring et al., 2011). In the current sample, the higher-order PTQ scale also showed excellent internal consistency (*α* = 0.95).

#### Ruminative Responses Scale (RRS, (Treynor et al., 2003))

Brooding rumination was assessed using the 5-item Brooding subscale of the Ruminative Responses Scale (RRS-B), which is considered the maladaptive form of rumination and has been implicated in the development of psychiatric disorders, such as depressive symptoms (Treynor et al., 2003). Participants responded using a 4-point Likert scale ranging from Almost Never (1) to Almost Always (4), rating how often they engage with an indicated thought or mental behaviour when down, sad, or depressed. Items included, for example, “Think ‘What am I doing to deserve this?’” or “Think about a recent situation, wishing it had gone better.” This scale has demonstrated acceptable reliability (*α* = 0.77) with a test-retest correlation of 0.62 in previous research (Treynor et al., 2003). In the current sample, the RRS-B showed good internal consistency (*α* = 0.81).

#### Anger Rumination Scale (ARS, (Sukhodolsky et al., 2001))

Anger rumination was assessed using the Anger Rumination Scale (ARS) (Sukhodolsky et al., 2001). The ARS comprises 19 items designed to measure the inclination to mentally engage with past episodes or feelings of anger. Participants are asked to rate each of the 19 items on a Likert scale ranging from Almost Never (1) to Almost Always (4), with higher scores indicating heightened anger rumination. Example items include statements like “Memories of even minor annoyances bother me for a while” and “I re-enact the anger episode in my mind after it has happened.” The ARS includes four subscales: angry afterthoughts, thoughts of revenge, angry memories, and understanding of causes. The ARS has previously exhibited adequate internal consistency, test-retest reliability, as well as both convergent and divergent validity (Sukhodolsky et al., 2001). In this study, the internal consistency was excellent (*α* = 0.93).

#### Screening for Anxiety and Depression in Tinnitus-Hyperacusis-Misophonia (SAD-T, (Aazh, Hayes, et al., 2022))

The 4-item Screening for Anxiety and Depression in Tinnitus-Hyperacusis-Misophonia (SAD-T) is a brief measure of anxiety and depression and has been validated in samples with auditory conditions (Aazh, Hayes, et al., 2022). A score of 4 or higher on this measure is considered the threshold for anxiety or depression symptoms. Participants are asked to rate how often they are bothered by the problems described in each item, with statements such as “Feeling nervous, anxious or on edge” and “Feeling down, depressed or hopeless.” In previous research, the SAD-T has shown good internal consistency (*α* = 0.91) and satisfactory ITC values between 0.76 and 0.84 (Aazh et al., 2024). In the current sample, the SAD-T showed excellent internal consistency (*α* = 0.90).

#### Sound Sensitivity Symptoms Questionnaire (SSSQ, (Aazh & Kula, Under review))

The Sound Sensitivity Symptoms Questionnaire (SSSQ) is a 6-item questionnaire designed to measure symptoms of abnormal sound intolerance, specifically hyperacusis and misophonia. Participants are asked to rate how often each of the statements apply to them, with scores on each item ranging from 0 to 3, corresponding to different frequency intervals (e.g. 0 to 1, 2 to 6,7 to 10, and 11 to 14 days). Example statements include “Pain in your ears when hearing certain loud sounds? Examples: loud music, sirens, motorcycles, building work, lawn mower, train stations” and “Fear that certain sounds may make your hearing and/or tinnitus worse?”. Total scores range from 0 to 18, with a total score of 4 considered to be the threshold for presence of sound sensitivity (Aazh et al., 2024). We included this measure to control for symptoms of hyperacusis, which has a high rate of co-occurrence with misophonia (Ahmmed & Vijayakumar, 2024; Andermane, Bauer, Sohoglu, et al., 2023). The previous version of SSSQ, which had 5 items, has shown good internal consistency (*α* = 0.87) as well as convergent and construct validity (Aazh et al., 2024). Here we utilized a new version with 6 items, which based on data from the authors showed a good Cronbach alpha estimate (*α* = 0.80), acceptable McDonald’s ω with values of 0.80, and test-retest reliability with an r-value of 0.81 well above Cohen’s suggested threshold for high correlation. In the current sample, the 6-item version showed acceptable internal consistency (*α* = 0.77) as well as when removing item-4 (which measures misophonia) from the scale (*α* = 0.71).

### Procedure

We first recruited participants through Prolific (www.prolific.com), an academic research-focused crowdsourcing platform. Eligibility criteria as screened using the demographic pre-screen feature on Prolific included age of 18-45, since individuals within this age range generally have similar audiological profiles, allowing control for hearing impairment/other periphery issues (Lee et al., 2012); no hearing loss or hearing difficulties; absence of cognitive impairment or dementia, normal or corrected-to-normal vision; fluency in English, and willingness to be exposed to distressing stimuli, considering the graphic nature of images included in the MAFT task. In addition, although no formal screening was used for comorbid psychiatric diagnosis, this was included as an exclusion criterion in the study advert and consent form (phrased as “No current psychiatric diagnosis or cognitive impairment” in the eligibility section).

Eligible participants based on Prolific pre-screen filter were shown a study advert briefly describing the procedures and also stating the compensation amount of $6.73 total USD (∼ €6.56 or ∼ £5.51). This compensation amount is equivalent to an estimated rate of $9.58 per hour (∼ €9.34 or ∼ £7.85) based on the median completion time of 42.06 min. Two participants, recruited from Prolific during the pilot stage of the study, were compensated in the approximate GBP equivalent to 6.73 USD based on the conversion rate at the time of the study.

After recruitment of participants via Prolific, we initiated an email campaign targeting newsletters (mentioned in participants section) aimed at individuals interested in volunteering for misophonia research. Those potential participants were sent the same study description and inclusion/ exclusion criteria used in the Prolific advert. Those who expressed interest were directed to a Qualtrics pre-screen survey to assess the same criteria (with identical wording of questions used to filter Prolific participants). Volunteer participants recruited from newsletters were not compensated for their time and were made aware of this before participation. This limitation arose from the requirement to exchange identifiable information, such as PayPal account details, which posed privacy and confidentiality concerns.

All participants were directed via URL to Qualtrics (www.qualtrics.com) to provide informed consent. This involved a description of the nature of images they might encounter during the MAFT task and participants were required to indicate their willingness to proceed knowing they would be exposed to such images. Upon providing consent, participants were redirected from Qualtrics to complete the Memory and Affective Flexibility Task via Inquisit Web version 6.6.1 (www.millisecond.com).

After completing the MAFT, participants were directed back to Qualtrics, where they re-entered their subject ID (Prolific-generated for Prolific participants, and Qualtrics-generated for volunteers), reported their age and gender, and then completed a battery of questionnaires (e.g., S-Five, rumination questionnaires, including the ARS, PTQ, RRS-B, SSSQ, SAD-T, and DFlex). Seven attention checks were included in the questionnaire battery, with one per questionnaire, in an Instructional Manipulation Checks format (e.g. “Please select ‘Often’ for this question”).

The entire study, including both MAFT and all questionnaires, took an estimated 40 to 45 mins, with a median study completion time of ∼42 mins. Upon adequate completion of the battery, Prolific-recruited participants were provided with a completion code and instructed to return to Prolific to receive compensation through the platform. Participants failing two or more attention checks received a custom code indicating failure and would have been requested to return their submission without compensation; however, no participants failed based on this criterion (**Figure 3**). Volunteer participants were not redirected upon completion, and the study concluded once they completed all components.

**Figure 3.**
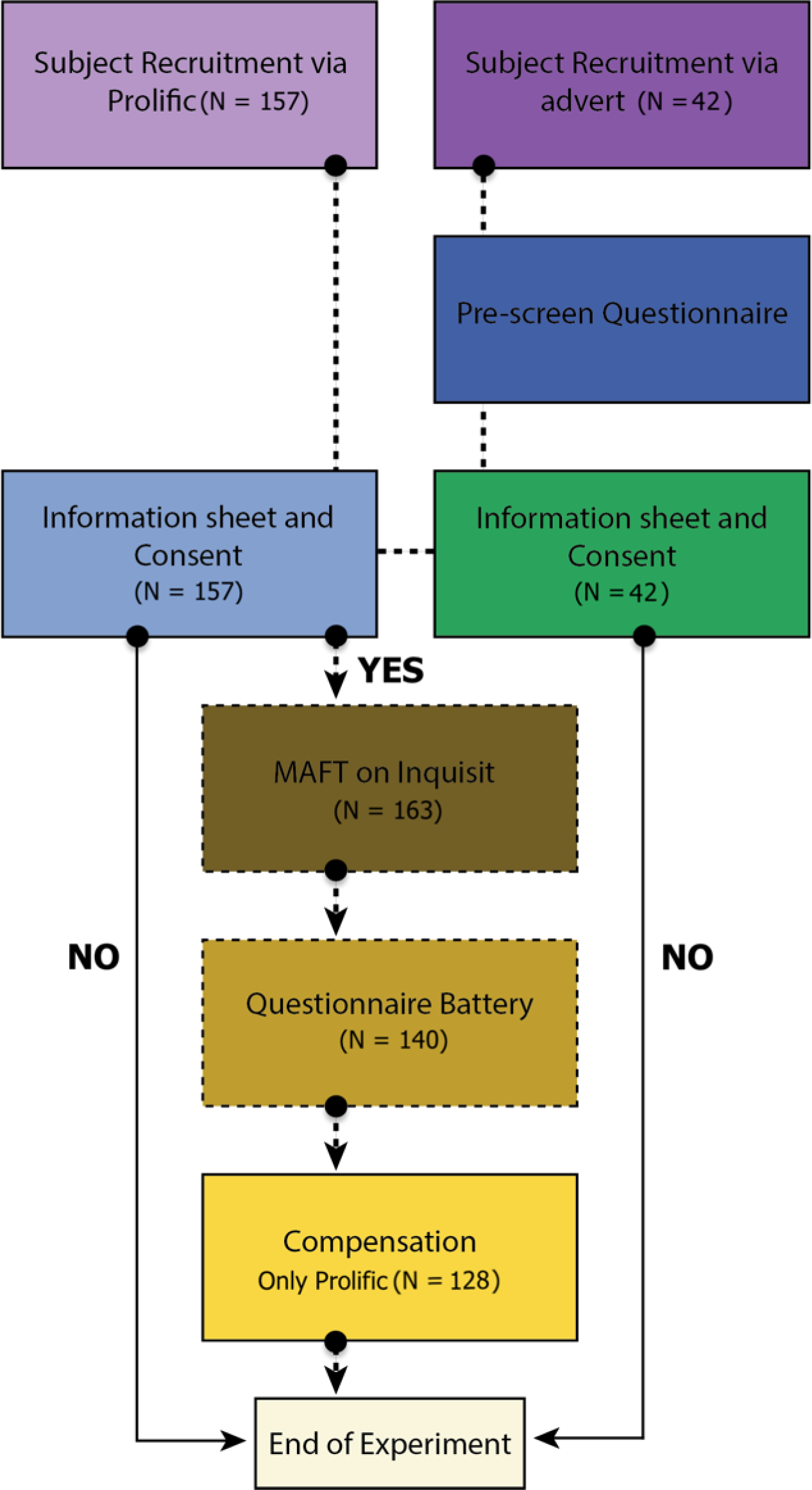
Diagram of online experiment progression. The diagram illustrates the study procedure, detailing each step from initiation to completion, along with the number of participants remaining at each stage. 199 subjects were initially recruited for the study. Of 157 Prolific participants, 142 attempted the MAFT, while 21 of 42 volunteers did so. From 163 total participants, exclusions included incomplete MAFT (N = 20), multiple attempts (N = 1), and incomplete questionnaires despite completing MAFT (N = 2), leaving N = 140. The final analysis included 140 participants who successfully completed the study. Only participants providing consent began the experiment. If the participants did not read the information sheet or did not consent, they were debriefed and directed to the end of the experiment.

Upon completion, Prolific-recruited participants’ self-reported demographic data became available for download via Prolific, with optional data first language, nationality, country of birth, student status, and employment status. This data was matched to participants’ Inquisit and Qualtrics data using Prolific ID; however, demographic data was not used for analysis purposes in the current study.

A total of 199 participants were recruited for the study, of whom 163 completed the MAFT: 142 from Prolific and 21 from outreach to newsletters. We excluded participants who did not complete the MAFT in full (*N* = 20) or attempted the MAFT task multiple times before completion (*N* = 1), or did not complete the Qualtrics questionnaires in full despite completing the MAFT (*N* = 2), leaving a final sample size of 140 (**Figure 3**). None of those participants failed two or more of the seven attention checks.

### Independent and Dependant variables

Our key independent variables included MAFT switch accuracy and switch RT indices, along measures of cognitive flexibility (DFlex), perseverative thinking (PTQ), brooding (RRS-B), and anger rumination (ARS). The dependent variables consisted of the S-Five total score and its five subscales. Possible confounding variables examined in the study were the total scores of the SSSQ and SAD-T, as well as gender.

The rationale behind controlling for hyperacusis as confounding variable was driven by the heightened SSSQ scores, suggestive of hyperacusis, in participants with significant misophonia. This pattern persisted even after excluding the misophonia-specific item (item 4). This is consistent with previous findings of heightened comorbidity with audiological disorders like tinnitus and hyperacusis (Aazh, Erfanian, et al., 2022). However, misophonia can also occur without audiological abnormalities (Swedo et al., 2022). Thus, where warranted, we include additional tests for all primary hypotheses with the SSSQ as a covariate in order to control for hyperacusis.

Evidence suggests that there is a high prevalence of depression and anxiety in individuals with misophonia (Aazh, Erfanian, et al., 2022; Danesh & Aazh, 2020; Erfanian et al., 2018; Erfanian et al., 2019; Erfanian & Rouw, 2018; Jager et al., 2020; McKay et al., 2018; Rouw & Erfanian, 2018; Siepsiak et al., 2020; Wu et al., 2014). Hence, we also control for depression and anxiety by incorporating the SAD-T score as a covariate in each regression model.

### Data Analytic Plan

#### Pre-processing

We pre-processed MAFT data using Python (Foundation, 2022), consistent with the methods of (Allen et al., Under review). All further statistical analyses were conducted using R version 4.3.1 (Team, 2023) and MATLAB 2019b (MATLAB, 2019) with an alpha set at 0.05. Bivariate Pearson correlations were calculated for all measures to assess overlap and correlations between variables.

Implausibly fast responses (< 200 ms) and missed responses (> 4500 ms), where participants were only allowed to register a response within the 4500 ms trial period, were excluded. Following the approach outlined in (Allen et al., Under review), trial-level outlier RTs with Z-scores > |3| were trimmed. Participants with a switch omission rate exceeding 0.5 were also excluded from analysis.

Across other measures, we reviewed outliers > 3 SD. These outliers (S-Five: two participants > 3 SD above the mean total score; ARS: one participant > 3 SD above the mean, and one < 3 SD below the mean) were retained in the data as they likely reflected natural variation, not measurement error. However, primary analyses were conducted both with and without outliers for comparison purposes to ensure consistency of findings.

Participants were categorized as experiencing clinical misophonia using a threshold score of 87 indicating *significant* misophonia (Vitoratou et al., 2023). Group differences were analysed using Χ² tests for nominal variables and Welch t-tests for continuous variables. Due to a significant difference in gender ratio between groups, primary analyses were repeated with gender as a covariate to ensure consistency.

#### Model specification (Multiple Linear Modelling)

To address the primary research questions, a series of Multiple Linear Models (MLMs) were conducted to examine the relationships between MAFT performance (switch accuracy and switch RT) as independent variables and misophonia severity, using the S-Five total score and its five subscales: externalizing, internalizing impact, threat, and outburst, as dependant variables. Separate parallel regression models were performed to investigate associations between cognitive flexibility, and rumination including the PTQ, RRS-B, and ARS as independent variables, misophonia severity and its subscales as the dependent variables. This resulted in 36 regression models. Gender, depression, anxiety and hyperacusis were included as covariates in models predicting S-Five scores to control for potential confounding effects. As no significant correlations were found between MAFT indices and the SAD-T or SSSQ measures, these variables were excluded as covariates in analyses of the relationships between switch accuracy and switch RT and S-Five scores.

Subsequently, we conducted independent regression models to examine the relationships between cognitive flexibility and rumination, including the PTQ, RRS-B, and ARS (dependent variables), resulting in a total of 3 models, while controlling for covariates.

For the mediation analyses exploring the interaction between rumination and cognitive factors in predicting misophonia severity, DFlex, ARS, RRS-B, and PTQ were included as independent variables, with the S-Five total score as the dependent variable. The same covariates were retained for consistency across analytical frameworks (3 models).

## Results

### Correlation between study variables

To establish the linearity between all pairs of variables, including independent, mediating and dependant variables, Pearson correlation coefficients were calculated. The variables analysed included the MAFT affective inflexibility indices including switch accuracy and switch RT, self-reported cognitive inflexibility measured by the DFlex, three rumination measures (PTQ, ARS, and RRS-B), misophonia severity as assessed by the S-Five total score and its subscales (externalizing, internalizing, impact, threat, and outburst), anxiety and depression symptoms assessed using the SAD-T, and hyperacusis measured by the SSSQ (**Figure 4**).

**Figure 4.**
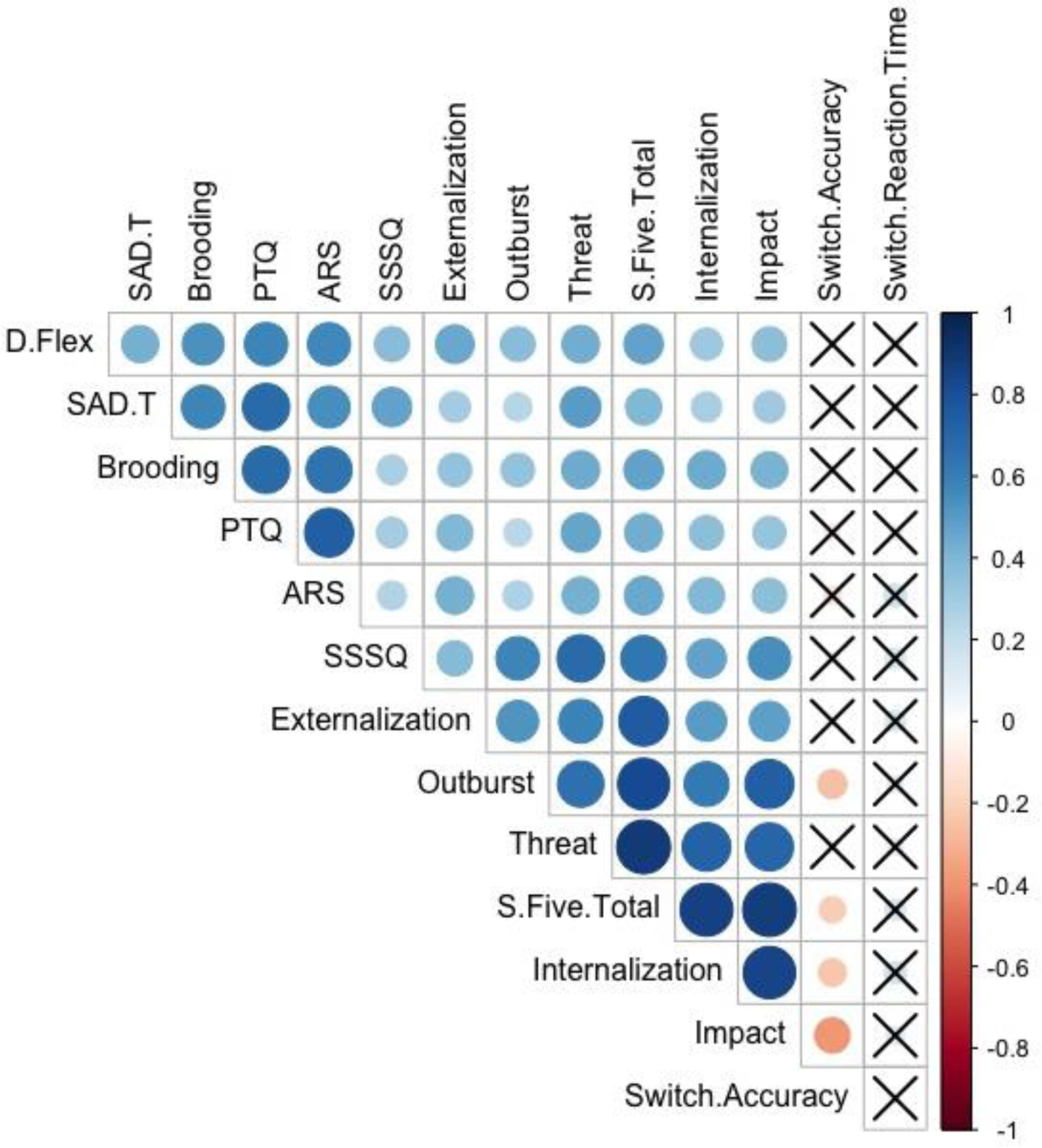
Correlation matrix of variables. The correlation matrix displays Pearson correlation coefficients among all study variables, with relationships represented by colour coding and size scaling. The size and colour gradient of the circles indicate the strength of the correlations, with warm colours (red) denoting negative correlations and cool colours (blue) denoting positive correlations. Circles marked with crosses represent non-significant correlations (p ≥ 0.05).

### Switch accuracy but not switch RT is associated with S-Five total, internalization, impact and outburst factors

Initially, we investigated the relationship of affective flexibility, as measured using switch accuracy and switch RT of the MAFT, with the severity of misophonia. As illustrated in **Figure 5**, switch accuracy on the MAFT demonstrated a significant inverse relationship with overall misophonia severity (total score; β = −0.21, t = −2.56, *p* = 0.01, as well as the internalizing (β = −0.24, t = −2.87, *p* < 0.01), impact (β = −0.38, t = −4.86, *p* < 0.001), and outburst domains (β = −0.26, t = −3.16, *p* < 0.01), but not the threat (β = −0.04, t = −0.48, *p* = 0.63) and externalizing (β = −0.06, t = −0.70, *p* = 0.49) domains. After excluding the two participants who scored > 3 SD above the mean S-Five total score, the relationship between switch accuracy and S-Five total score remained significant (β = −0.24, t = −2.94, *p* < 0.01). Furthermore, the relationship between switch accuracy and misophonia severity was robust after controlling for gender (β = −0.24, t = −2.88, *p* < 0.01).

**Figure 5.**
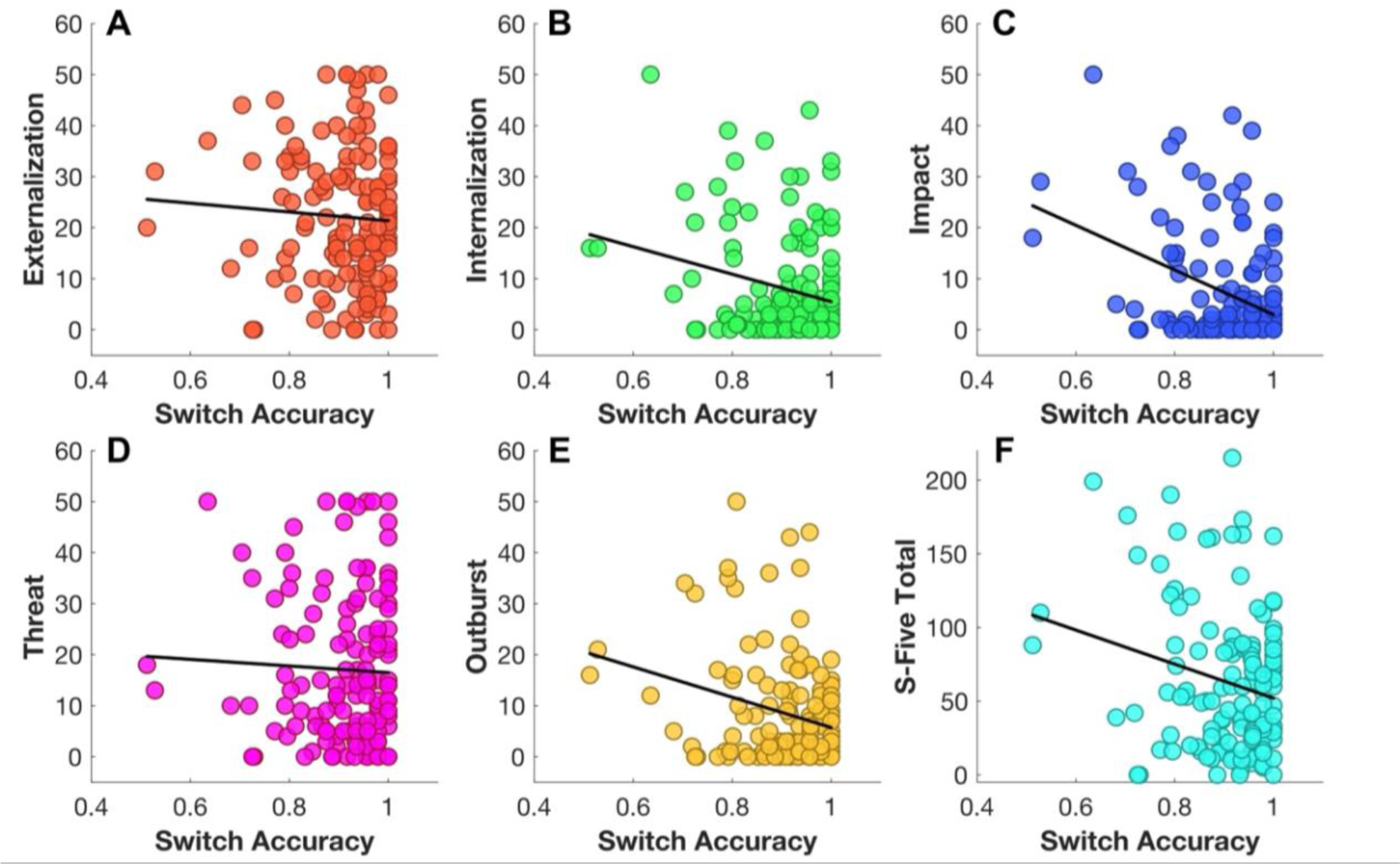
Scatterplot of bivariate analyses. The scatterplots demonstrate the bivariate relationships between MAFT switch accuracy as a proxy of affective flexibility and **A)** externalization, **B)** internalization, **C)** impact, **D)** threat, **E)** outburst as S-Five subscales and **F)** S-Five Total, reflecting misophonia total severity. The circles represent individual data points, while the diagonal black lines indicate the line of best fit.

As shown in **Figure 4**, switch accuracy was not significantly correlated with SAD-T score (β = −0.04, t = −0.49, *p* = 0.62) nor with the SSSQ (misophonia question excluded), (β = −0.07, t= −0.79, *p* = 0.43), indicating that the relationship between switch accuracy and misophonia severity may be independent of comorbid anxiety, depression, or hyperacusis.

Given the absence of significant correlations between the switch RT index and the other variables of interest, as determined by Pearson correlation analysis, switch RT was excluded from subsequent multivariate analyses. Although not significant, our findings are consistent with those of (Allen et al., Under review), reporting that RT indices were highly correlated across all MAFT summary metrics of affective flexibility. This pattern suggests that RT scores may primarily reflect individual differences in processing or motor response speed rather than specific effects of task conditions (Allen et al., Under review).

### Participants with significant misophonia score worse for switch accuracy on the MAFT but do not demonstrate a slower switch RT

Given that a substantial proportion of our sample scored below the threshold for significant misophonia (*N* = 105), participants were divided into two groups based on a threshold score of 87. Switch accuracy and RT were then compared between those with lower misophonia severity (<87) and those with higher severity (>87) (significant misophonia). As shown in **Figure 6**, we found a significant between-group difference in mean switch accuracy, with individuals classified as experiencing significant misophonia showing worse accuracy on the MAFT switch trials, with a medium effect size (t = 2.61, *p* = 0.01, d = 0.67). No significant difference, however, was observed comparing the two groups’ switch RT data (t = −0.13, *p* = 0.90, d = −0.03), in line with our continuous-scale findings. This demonstrates that the relationship between switch accuracy and misophonia is also evident dichotomously when comparing groups based on the S-Five cutoff threshold.

**Figure 6.**
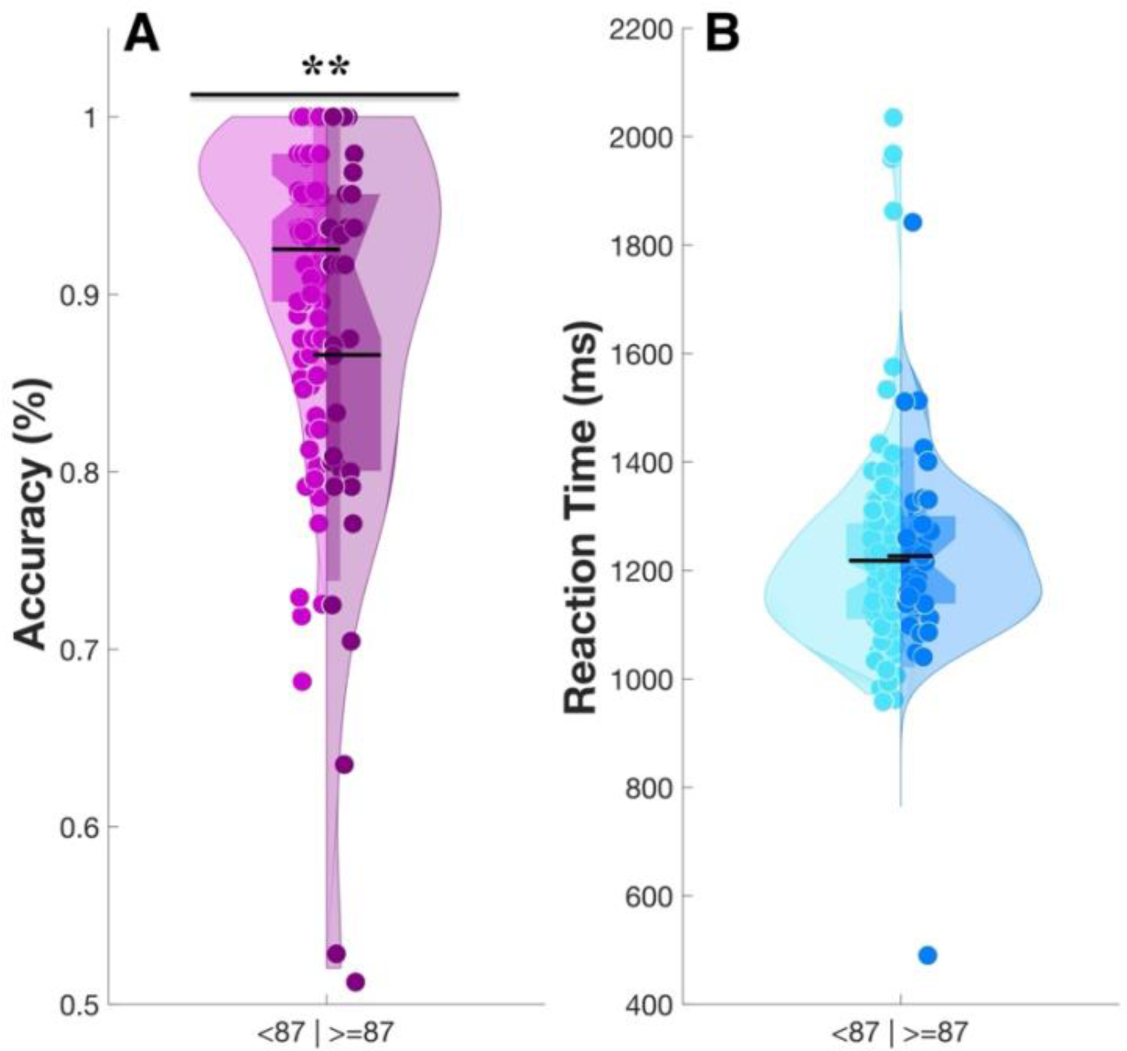
The violin plots of misophonia groups. The figures present the results of Welch t-test comparing between-group differences in MAFT affective flexibility performance, measured by switch accuracy **(A)** and switch RT **(B)**, between participants with significant misophonia (scores ≥ 87) and those without. The X-axis represents two groups: those scoring below and above 87, while the Y-axis displays accuracy as a percentage (%) for switch accuracy and switch RT (ms). Individual data points are represented by single circles. The bottom edge of the thick rectangle represents the 1st quartile (Q1, 0.25), while the top edge represents the 3rd quartile (Q3, 0.75). The lower end of the kernel density curve indicates the 1^st^ percentile (Q 0.01), and the upper end represents the 99^th^ percentile (Q 0.99). The bottom of the long bar illustrates the standard deviation, while the black horizontal line indicates the mean. The top asterisk denotes statistical significance between the groups. *p < 0.05, ** p < 0.01, and ***p < 0.001.

### S-Five total and all its subscales are associated with decrease in cognitive flexibility score on DFlex

To further explore whether impaired mental flexibility potentially extends beyond the behavioural task and can be extrapolated to broader contexts, we tested the replicability of Simner et al., (2021). As shown in **Figure 4**, Pearson correlation analysis established a significant moderate relationship between S-Five score (total and all 5 factors) and DFlex flexibility subscale score. Regression models confirmed the significance of cognitive inflexibility score as predictor of S-Five total: β = 0.26, t = 3.44, *p* < 0.001, externalizing: β = 0.38, t = 4.42, *p* < 0.001, impact: β = 0.19, t = 2.27, *p* = 0.02; threat: β = 0.15, t = 2.00, *p* < 0.05; outburst: β = 0.21, t = 2.53, *p* = 0.01). Controlling for gender, SSSQ, and SAD-T as covariates, however, DFlex score did not account for significant variance on the internalizing domain of the S-Five (β = 0.16, t = 1.78, *p* = 0.08). In light of these observations, we conducted additional analyses to explore the emergence of this pattern further. The model was re-run after excluding SAD-T, revealing a noteworthy, albeit marginal, association between DFlex scores and internalization (β = 0.178, t = 2.069, *p* = 0.04). Integrating these results, our findings indicate that misophonia is significantly associated with self-reported cognitive inflexibility on a continuous scale. These outcomes not only corroborate but also expand upon the findings of (Simner et al., 2021). Furthermore, it appears that the relationship between cognitive inflexibility and the internalization factor is predominantly accounted for by the co-occurrence of anxiety and depression, highlighting a potential mediating role for these affective states. Although this result holds considerable importance, it is noteworthy that staggering evidence show that depression and anxiety are primarily internalizing disorders which help to explain this connection (Doering et al., 2022; Kovacs & Devlin, 1998; Priest & Cobb, 2020; Tandon et al., 2009) (**Figure 7**).

**Figure 7.**
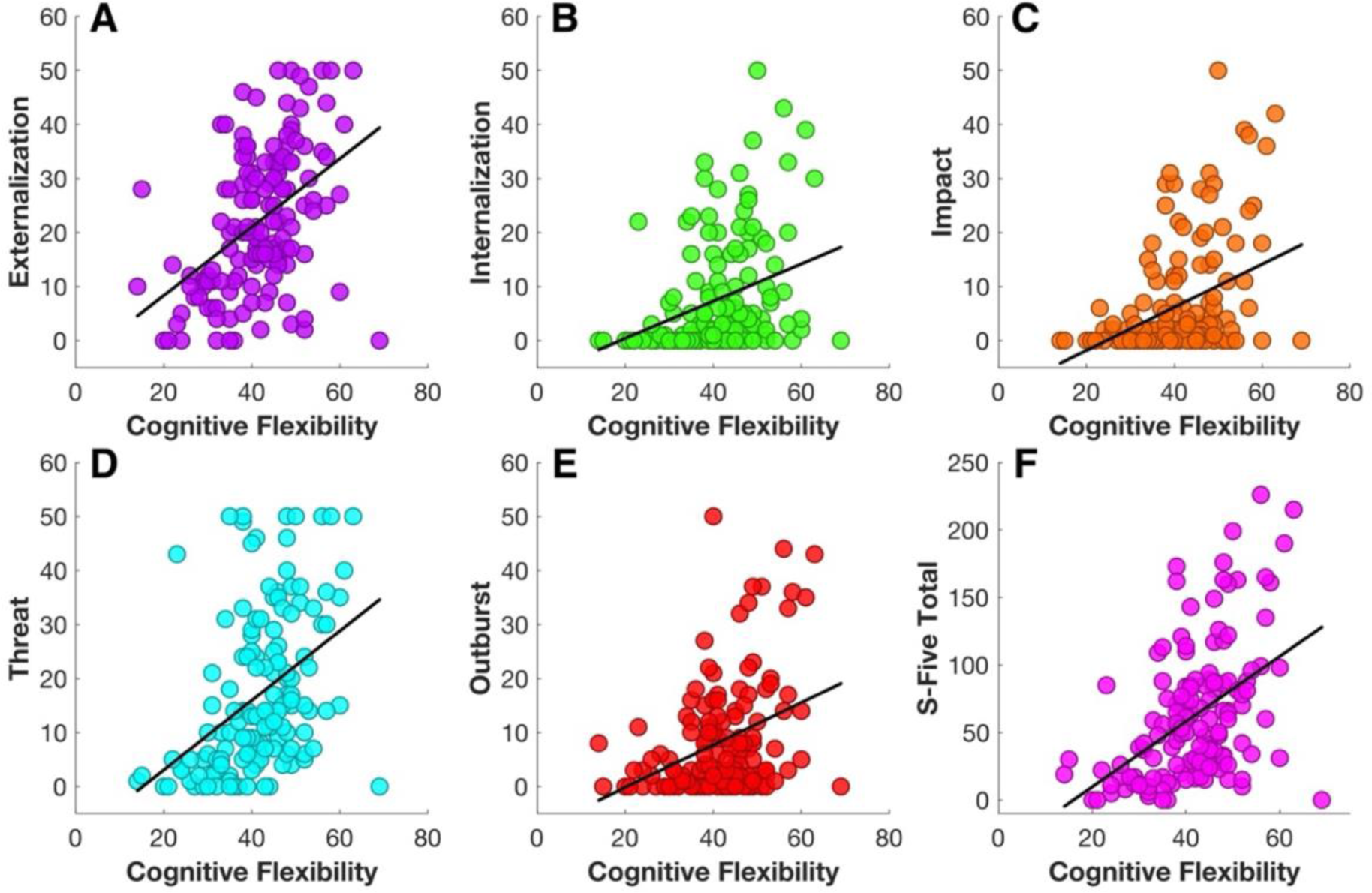
Scatterplot of bivariate analyses. The scatterplots show the bivariate relationships between DFlex representing cognitive flexibility and **A)** externalization, **B)** internalization, **C)** impact, **D)** threat, **E)** outburst as S-Five subscales and F) S-Five Total, reflecting misophonia total severity. The circles represent individual data points, whereas the diagonal black lines show the line of best fit.

### Switch accuracy or switch RT of MAFT are not associated with score on DFlex

Our findings indicated no significant correlation between affective flexibility (MAFT) and self-reported cognitive flexibility (switch accuracy: r = 0.02, *p* = 0.79; switch RT: r = 0.07, *p* = 0.43). These results suggest that while misophonia is linked to cognitive inflexibility, the observed relationships are likely independent of the mechanisms captured by switch accuracy-based measures of affective flexibility in the MAFT paradigm. These findings highlight the notion that task-based and questionnaire-based measures may overlap conceptually but likely represent distinct psychological constructs.

### Self-reported rumination is associated with total score of S-Five, and effects persist when controlling for gender, SAD-T and SSSQ

As shown by **Figure 4**, Pearson correlation analyses revealed moderate positive associations between all three forms of rumination and misophonia severity (including all S-Five subscales). We then used regression models to explore the relationship between three forms of rumination, perseverative thinking (PTQ), brooding (RRS-B), anger rumination (ARS), and misophonia severity while accounting for the influence of potential confounding variables: anxiety and depression, gender, and comorbid hyperacusis.

Using a separate regression equation for each rumination questionnaire (three total, shown in **Figure 8**), each subtype of rumination remained significant predictor of misophonia severity (measured by S-Five total) when controlling for SAD-T score, gender, and SSSQ score (PTQ: β = 0.36, t = 4.05, *p* < 0.001; ARS: β = 0.33, t =4.37, *p* < 0.001; RRS-B: β= 0.36, t = 4.65, *p* < 0.001), and also when excluding participants who reached outlier scores on the S-Five (PTQ: β = 0.35, t = 3.92, *p* < 0.001; ARS: β = 0.43, t = 4.30, *p* < 0.001; RRS-B: β = 0.32, t = 4.04, *p* < 0.001) or the one ARS outlier (β = 0.48, t = 4.81, *p* < 0.001).

**Figure 8.**
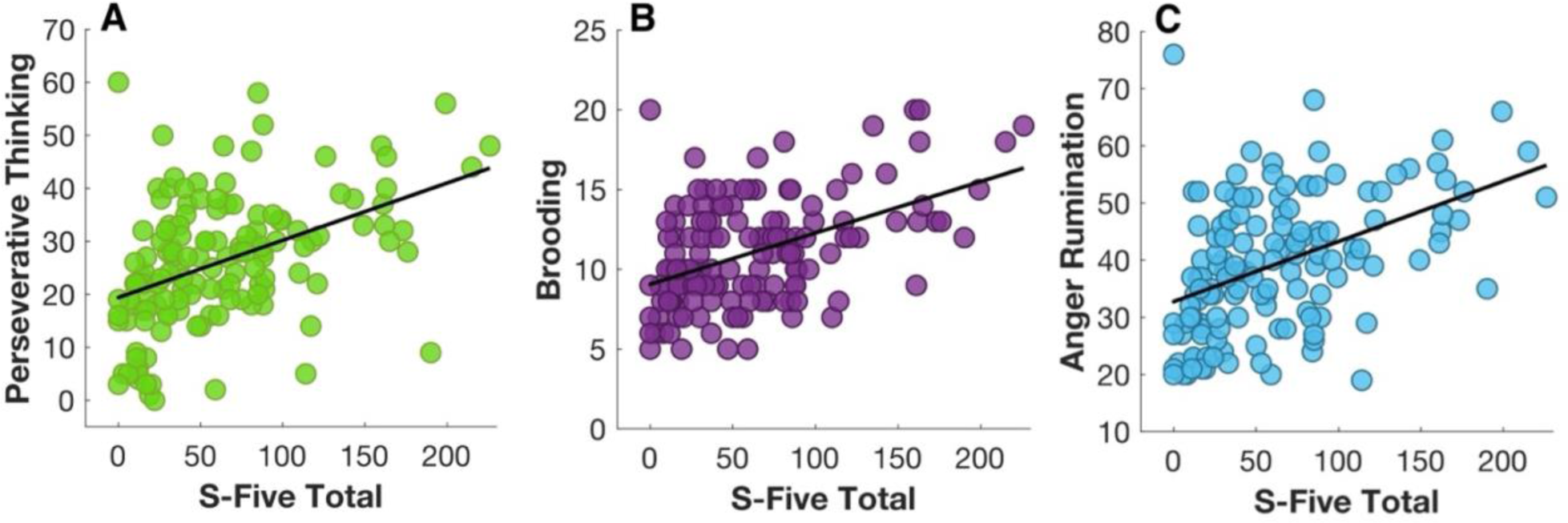
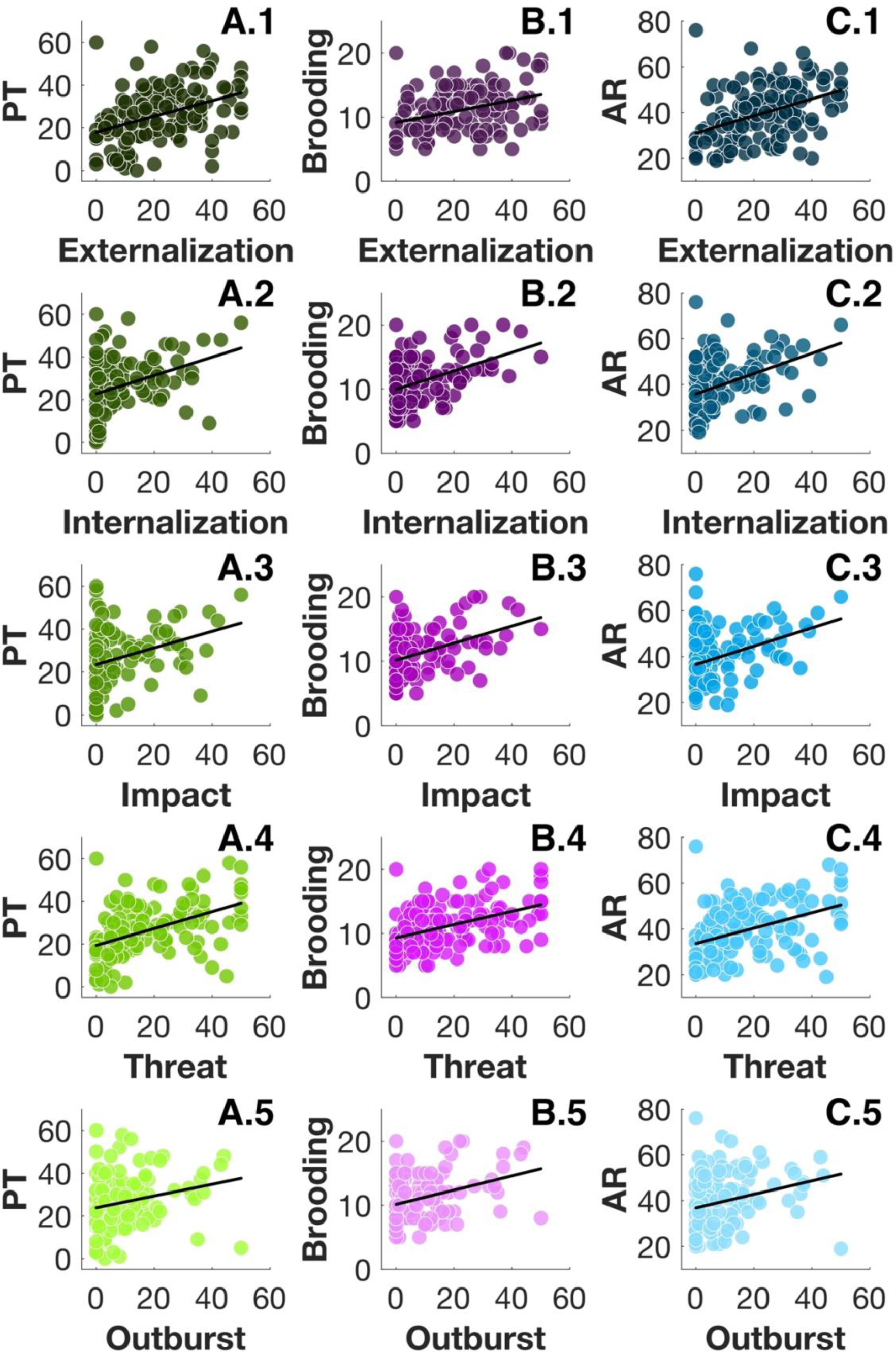
The scatterplots illustrate the bivariate relationships between S-Five Total, reflecting misophonia severity (**A, B, and C**) and its subscales: **1)** externalization, **2)** internalization, **3)** impact, **4**) threat, and **5)** outburst with **A.1 to A.5)** perseverative thinking (PT), **B.1 to B.5)** rumination, and **C.1 to C.5)** anger rumination (AR). The circles represent individual data points, while the diagonal black lines indicate the line of best fit.

In more details, perseverative thinking was associated with externalization (β = 0.42, t = 3.81, *p* < 0.001), internalization (β = 0.30, t = 3.55, *p* < 0.001), impact ( β = 0.23, t = 2.79, *p* = 0.006), threat (β = 0.33, t = 3.54, *p* < 0.001), and outburst (β = 0.14, t = 1.76, *p* = 0.08). We observed similar pattern for brooding and externalization (β = 1.02, t = 2.77, *p* = 0.006); internalization (β = 1.29, t = 4.98, *p* < 0.001), impact (β = 1.099, t = 4.32, *p* < 0.001); threat (β = 1.002, t = 3.31, *p* = 0.001); and outburst (β = 0.94, t = 3.75, *p* < 0.001). Interestingly, the same pattern continues in anger rumination with externalization (β = 0.43, t = 4.33, *p* <001); internalization (β = 0.31, t = 4.15, p < 0.001); impact (β = 0.25, t = 3.41, *p* <0.001); threat (β = 0.26, t = 3.13, *p* <0.001); and outburst (β = 0.17, t = 2.35, *p* = 0.02) (**Figure 8**).

These results show that rumination, not limited to one subtype, is positively associated with misophonia severity and all five dimensions of misophonia. Although both rumination and misophonia are associated with anxiety, depression and hyperacusis (**Figure 4**), these findings imply that rumination explains variance in misophonia over and above the variance explained by anxiety/depression, hyperacusis and gender.

### Self-reported rumination is unrelated to MAFT switch accuracy or switch RT, but linked to DFlex

As shown in **Figure 4**, Pearson correlation analysis established no significant relationship between switch accuracy any of the three rumination subscales: ARS: r = −0.11, *p* = 0.20; PTQ: r = −0.07, *p* = 0.40; RRS-B: r = −0.02, *p* = 0.83. The same was true for switch RT, which also showed no significant relationship with the rumination questionnaires based on correlation analyses: PTQ: r = 0.06, *p* = 0.48; RRS-B: r = 0.04, *p* = 0.67; ARS r = 0.16, *p* = 0.06.

However, the correlation analyses yielded significant relationships between DFlex score and all three forms of rumination and this was confirmed with regression analyses (ARS: β = 0.42, t = 5.57, *p* < 0.001; PTQ: β = 0.39, t = 6.04, *p* < 0.001; and RRS-B: β = 0.35, t = 4.72, *p* < 0.001), when controlling for the confounding factors. The observed relationship between cognitive flexibility and rumination, despite the MAFT indices showing no significant association with rumination, points to the notion that cognitive and affective flexibility, while closely interconnected, may represent distinct constructs (**Figure 9**).

**Figure 9.**
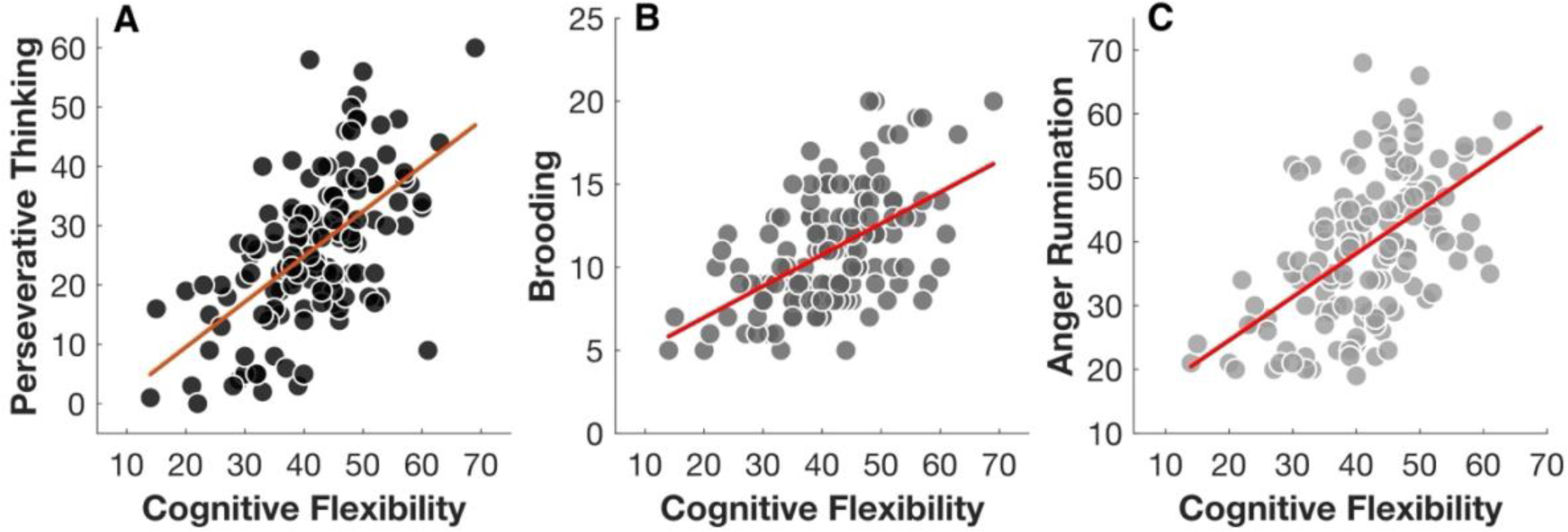
The scatterplots show the bivariate relationships between DFlex, representative of cognitive flexibility and **A)** perseverative thinking, **B)** brooding, and **C)** anger rumination. The circles represent individual data points, while the diagonal red lines indicate the line of best fit.

Since significant associations were observed between the S-Five total score and all three types of rumination (perseverative thinking, brooding, and anger rumination) alongside cognitive flexibility, it is reasonable to argue that the interplay between rumination and cognitive flexibility may better explain the variations in S-Five total. To explore this interaction, separate regression models were applied with PTQ, RRS-B, ARS and DFlex as independent, and S-Five total as the dependant variables, while controlling for depression, anxiety and hyperacusis.

The analyses revealed nuanced interaction effects across the models. In Model 1 (*R^2^* = 0.298, *p* < 0.001), DFlex emerged as a negative moderator of the relationship between PTQ and misophonia severity, as evidenced by the interaction term coefficient (*b* = −0.022). This finding indicates that greater cognitive flexibility mitigates the adverse influence of perseverative thinking on misophonia symptoms. Similarly, in Model 2 (*R^2^* = 0.29, *p* < 0.001), the interaction term coefficient (*b* = −0.019) suggested that DFlex attenuated the effect of RRS-B on misophonia severity. In Model 3 (*R^2^* = 0.278, *p* < 0.001), the interaction term coefficient (*b* = −0.031) demonstrated a comparable moderating effect of DFlex on the relationship between ARS and misophonia severity. These findings point to the complex and context-dependent role of cognitive flexibility in moderating the effects of rumination on misophonia. In essence, cognitive flexibility may buffer the detrimental impact of certain types of rumination, highlighting the need for a nuanced understanding of its interaction with cognitive and emotional processes in the context of misophonia.

## Discussion

In the current work, we extend previous research on cognitive inflexibility in misophonia in several ways. First, we conjointly considered a self-report measure of broad cognitive flexibility, together with a behavioural measure of affective flexibility. Second, we considered how misophonia and cognitive inflexibility related to multiple forms of rumination. Third, throughout our analyses, we considered multiple dimensions of misophonia symptoms. Consistent with hypotheses, we demonstrated that affective flexibility, limited to switch accuracy, was inversely associated with misophonia severity and internalization, impact and outburst factors. We also observed that cognitive inflexibility was significantly associated with the severity of misophonia and these findings generalized across misophonia subscales, except for internalization. Consistent with previous work, we found that cognitive flexibility and affective flexibility were separable, with no significant correlation between these indices. Furthermore, our findings revealed that various types of rumination, including perseverative thinking, brooding, and anger-related rumination, were associated with misophonia severity and cognitive inflexibility but not affective inflexibility.

Taken together, these findings imply that the mechanisms of misophonia may not be specific to sound triggers. Instead, misophonia appears to involve a complex interplay of cognitive and emotional processes, which may contribute to the heightened sensitivity and exaggerated responsivity to specific auditory stimuli. That is, we observed reduced task-switching accuracy when faced with emotion-evoking stimuli that were unrelated to auditory triggers, which suggests the broader cognitive affective contributors to this condition, beyond reactions specifically linked to misophonic triggers. Findings also highlight the potential role of maladaptive emotional regulation strategies, such as rumination, in the pathology of misophonia. Overall, our findings emphasize the need to consider cognitive and affective processes in understanding this condition.

### Switching to affective stimuli: Greater challenges linked to increased misophonia symptom severity

To provide evidence of affective flexibility deficits in individuals with misophonia, we employed a newly developed task designed to capture affective switching. This task allowed us to explore the relationship between two performance indices, switch accuracy and switch RT, and misophonia severity. As expected, overall misophonia severity and affective flexibility, limited to switch accuracy, were inversely associated. RT performance was not related to misophonia, and this is consistent with early evidence that RTs on this task are not specific to affective flexibility but tend to reflect a more general motor response rate across conditions (Allen et al., Under review). The diminished affective flexibility may hamper the ability to effectively switch responding to demands in the face of emotional stimuli. This finding is in line with previous studies which highlight the role of emotion regulation in misophonia (Bagrowska et al., 2022; Barahmand et al., 2023; Cassiello-Robbins et al., 2020; Dixon et al., 2024; Guetta et al., 2022).

Similar patterns have been revealed in individuals with high autistic traits, overlapping with misophonia symptoms (Ertuerk et al., 2023; Rinaldi et al., 2023), who show greater switch costs (worse accuracy), suggesting potential links between cognitive inflexibility and the expression of autistic traits (Zhou et al., 2024). Two broad, mutually inclusive classes of explanation can be distinguished to account for the increased switch costs observed in individuals with misophonia. The Task-Set Reconfiguration (TSR) Theory (Rogers & Monsell, 1995) posits that switch costs arise from the cognitive effort required to reconfigure a new task set during switch trials, a process reliant on endogenous, top-down control mechanisms (Zhou et al., 2024). Evidence suggests that cognitive control, a mechanism that facilitates task reconfiguration, is impaired in individuals with misophonia when confronted with trigger sounds that are necessary for reconfiguring the task (Daniels et al., 2020). In addition, effective cognitive control is crucial for deploying response inhibition when needed, such as restraining automatic but contextually inappropriate behaviours, like anger outbursts in response to trigger sounds. Altered response inhibition related to misophonia also has been observed using the stop-signal task, revealing a bias toward longer stop-signal delays (Eijsker et al., 2019). According to this framework, the larger emotion switch costs observed individuals with misophonia may reflect difficulties in reconfiguring the emotional task set.

Alternatively, the Task-Set Inertia (TSI) Theory (Allport & Wylie, 1999; Meiran, 2008; Waszak et al., 2003; Wylie & Allport, 2000) asserts that switch costs arise from interference caused by residual activation of previous task sets. From this perspective, the larger emotion switch costs in misophonic individuals may indicate greater susceptibility to interference from prior task sets when transitioning to an emotional task. Both task-set inertia and task-set reconfiguration are thought to jointly contribute to the increased emotion switch costs (Vandierendonck et al., 2010).

The reduced affective flexibility in misophonia can also be better understood by the neural correlates of misophonia. Inhibition is essential for set-switching within affective flexibility, facilitating the shift between behaviour, emotion or cognitive process or task sets by suppressing competing or irrelevant ones and minimizing interference during the transition (Arbuthnott & Frank, 2000). Elevated orbitofrontal cortex (OFC) activity has been documented during successful, as opposed to failed, behavioural inhibition, highlighting its role in regulatory control processes (Deng et al., 2017). In individuals with misophonia, the OFC exhibits hyperactivation specifically in response to trigger stimuli. This heightened activity suggests the critical role of OFC in facilitating functional synchronization among brain regions, potentially influencing the misophonic ‘response’ as reflected in behavioural task performance (Cerliani & Rouw, 2020).

### Heightened misophonia severity links to reduced cognitive flexibility

Confirming our behavioural findings and replicating the results of (Simner et al., 2021), individuals with more severe misophonia showed reduced cognitive flexibility. These findings imply a broader contextual rigidity in misophonia, extending beyond specific behavioural responses. Such rigidity has been observed in other overlapping psychiatric and neurodevelopmental disorders (Geurts et al., 2009; Gruner & Pittenger, 2017; Miles et al., 2020; Murphy et al., 2012) with high co-morbidity with misophonia (Erfanian et al., 2019; Erfanian & Rouw, 2018; Jager et al., 2020; Rouw & Erfanian, 2018).

Cognitive inflexibility in misophonia may account for heightened sensitivity to norm violations in uncontrollable conditions (Banker et al., 2022), reflecting a diminished ability to adapt to environmental deviations or unpredictability (Miyake et al., 2000) and to flexibly adjust cognitive strategies to manage change (Han et al., 2012). Similar to individuals with autism, those with cognitive inflexibility often resist change and struggle to shift their mental frameworks or expectations, leading to a rigid adherence to established norms (Lacroix et al., 2024). Consequently, when faced with uncontrollable situations that deviate from these norms, their difficulty in adapting results in heightened sensitivity and an amplified reaction to perceived violations (Norena, 2024). This rigidity intensifies discomfort and limits their ability to contextualize or tolerate variability

Although mental rigidity, attributed to misophonia, is likely a key explanation for the tendency to over-attend or hyperfocus in response to misophonic triggers (Edelstein et al., 2013; Jager et al., 2020; Simner et al., 2021), it may also represent an underlying pathological mechanism that contributes to broader perceptual and behavioural deficits. To advance the field and explore whether the behavioural symptoms are underpinned by the rigidity or rigidity should be considered *per se* as a symptom, experimental measures must evolve to reflect mechanistic models of flexibility deficits.

### The severity of misophonia is linked to elevated rumination tendencies

Building on the robust evidence linking cognitive flexibility and rumination across various psychopathological conditions, we delved into the potential association of misophonia with three forms of rumination (Altan-Atalay et al., 2022; Davis & Nolen-Hoeksema, 2000; Genet et al., 2013; Lei et al., 2022; Miles et al., 2023; Owens & Derakshan, 2013). All three forms of rumination were moderately associated with misophonia severity, and these relationships remained significant even after controlling for depression, anxiety and hyperacusis. This finding implies that the relationship between rumination and misophonia severity cannot be explained by the presence of these comorbid symptoms. Although rumination is a hallmark feature of depressive and anxiety disorders (Ruscio et al., 2015), both of which frequently co-occur with misophonia (Erfanian et al., 2018; Erfanian et al., 2019; Erfanian & Rouw, 2018; Jager et al., 2020; Rosenthal et al., 2022; Rouw & Erfanian, 2018), our findings emphasize the unique connection between rumination and misophonia severity.

Anecdotal reports, consistent with our findings, indicate that individuals with misophonia employ internal coping strategies. These strategies involve cognitive processing of trigger characteristics, anger rumination, and contemplation of emotional responses associated with misophonia, aimed at avoiding or escaping triggers (Dozier & Mitchell, 2023).

Yet, these findings should be interpreted in the context of cognitive flexibility, especially given its association with all forms of rumination within this cohort. First, the inability to mentally switch between cognitive sets may lead to perseveration (Altan-Atalay et al., 2022; Beckwé et al., 2014; Nolen-Hoeksema, 2000; Watkins & Brown, 2002). Second, similar to mood disorders, the ruminative tendencies observed in misophonia could reflect disturbances in cognitive switching processes (Piguet et al., 2010), leading to maladaptive emotion regulation strategies that are less effective in the moment (Barahmand et al., 2023; Cassiello-Robbins et al., 2020; Dixon et al., 2024; Erfanian et al., 2018; Guetta et al., 2022; Rinaldi et al., 2023; Spencer et al., 2023).

It is important to note that we were not able to disentangle the conjoint effects of cognitive flexibility vs. rumination in relation to misophonia. All forms of rumination and cognitive inflexibility overlap in their relationship with misophonia. Previous research supports the bidirectional effects of cognitive flexibility and rumination (De Lissnyder et al., 2012). For instance, in the context of AN, which shows high comorbidity with misophonia, evidence suggests that eating disorder-specific rumination acts as a mediator between subjective cognitive flexibility and eating disorder symptoms (Miles et al., 2023).

All in all, as we measured multiple forms of rumination and focused on general tendencies of responding to distress rather than misophonia-specific experiences, our findings demonstrate that a tendency toward perseverative thought in misophonia extends beyond trigger-related situations. Fortunately, perseverative thought such as rumination can be targeted in treatment through therapy modalities such as CBT (Querstret & Cropley, 2013), which has shown promise in managing misophonia symptoms (I. J. Jager et al., 2020).

## Conclusion

This study provides novel evidence that the severity of misophonia is related to affective inflexibility, cognitive inflexibility, and perseverate thinking. Affective inflexibility was distinct to cognitive inflexibility and perseverative thinking, as no significant association was observed with measures of these constructs. These findings provide insights into the rich set of affective and cognitive dimensions involved in misophonia. The profile of affective and cognitive difficulties observed here are well-documented as transdiagnostic risk variables, consistent with the diagnostic overlap of misophonia with conditions such as ASD and OCD. By delineating these characteristics, the study contributes to reducing the risk of misdiagnosis and enhances our understanding of thee cognitive and affective processes involved in misophonia. Importantly, this research offers practical implications for clinical practice, encouraging a more targeted approach to treatment protocols that emphasise addressing the cognitive aspects of misophonia.

## Acknowledgement

This study was generously funded by the University of California, Berkeley Undergraduate Research Apprentice Program (URAP) and the SoQuiet Misophonia Student Research Grant (https://www.soquiet.org/grants). ME and HA received additional support from the Research and Development Fund provided by Hashir International Specialist Clinics & Research Institute for Misophonia, Tinnitus, and Hyperacusis Ltd.

## Author contribution

VB and ME conceptualized the study and developed the methodology with input from JDA. VB conducted the data collection, formal analysis and curated the data with supervision of SJ, JDA and ME. Software development and resources were managed by VB and JDA, while ME took the lead on validation and visualization. The original draft of the manuscript was written by VB and ME and editing carried out collaboratively. HA, SLJ and ME supervised the project. SLJ and ME oversaw the project administration and supported funding acquisition efforts.

## Data availability

All raw and processed data are available at [http://osf.io/KDRCE].

